# Dual Inference Routes for Natural Language Hypothesis Generation in Multimodal Drug Discovery: An Early Experimentation Study

**DOI:** 10.1101/2025.09.02.673688

**Authors:** Ren Wu, Hiroshi Matsuno, Qi-Wei Ge

**Affiliations:** Shunan University, Faculty of Information Science, Shunan, 745-8566, Japan; Yamaguchi University, Graduate School of Sciences and Technology for Innovation, Yamaguchi, 753-8511, Japan; Yamaguchi University, Faculty of Education, Yamaguchi, 753-8511, Japan

## Abstract

This study presents early experimentation of a modular AI framework for multimodal drug discovery that integrates natural product-based therapies with modern pharmaceuticals. The system combines structured biomedical data, knowledge graphs, and large language models (LLMs) to generate explicit natural language hypotheses. The architecture has four phases: data aggregation, hypothesis generation, dynamic simulation, and *in silico* evaluation, and supports dual inference routes (compound → gene → disease/phenotype and disease/phenotype → gene → compound). As a case study, Phases 1 and 2 were applied to the Kampo formula *Shakuyaku-kanzo-to*, a typical example of a multicomponent and multi-target natural therapy. The framework originally arose from challenges in conventional filtering, where important but poorly annotated compounds were often overlooked. However, the focus has since shifted beyond filtering, toward uncovering hidden relationships across fragmented biomedical knowledge. This early implementation demonstrates the potential of natural language hypothesis generation to restructure fragmented knowledge into interpretable insights, providing a blueprint for future multimodal drug discovery.

## 1 Introduction

Conventional drug discovery has traditionally followed the *one-drug-one-target* paradigm, a longstanding cornerstone of Western medicine^1^. While this approach has yielded effective treatments for many diseases, it also shows fundamental limitations. These include notable side effects, limited efficacy in treating complex systemic conditions such as depression and chronic inflammation, and the high cost and frequent failure rates of modern drug development^2,3^.

In contrast, the principles of multicomponent therapeutic systems, such as Kampo, align well with modern network-based approaches that seek to understand the multiple actions of drugs through complex biological and chemical interaction networks^4^. In particular, network pharmacology is regarded as a useful framework for clarifying the multi-target mechanisms of therapies derived from natural products^5–8^. Although the individual constituents are sometimes perceived as *weaker*, their overall therapeutic effects may arise from the complex bioactivity of their numerous interacting components. This concept is well-supported by comprehensive reviews covering both systems-level mechanisms and modern biological approaches to pharmacology^9,10^, offering new opportunities when integrated with modern drug discovery.

This study proposes a multimodal pharmaceutical framework that integrates modern drugs with naturally derived compounds. By integrating empirical herbal knowledge with pathway-based mechanisms into a unified architecture, the system aims to facilitate transparent and systematic hypothesis generation for complex diseases.

We initially explored conventional data-driven filtering approaches, but encountered key limitations that underscore the risks of relying solely on structured databases. For instance, paeoniflorin, a well-known constituent of *Paeoniae Radix*, is often excluded from network-based analyses due to the scarcity of documented protein targets. This challenge of insufficient target information is a known issue, prompting researchers to employ computational discovery approaches even for such well-known compounds^11^. More critically, this data scarcity introduces a research bias, concentrating attention on a few well-documented molecules while leaving the broader chemical space of potentially bioactive constituents underexplored^12^. These limitations, symptomatic of broader challenges in data standardization and reliability in the field^13^, highlight the urgent need for novel frameworks that can overcome the constraints of conventional filtering.

To address this limitation, we combine large language models (LLMs) with structured biomedical databases (e.g., STITCH, ChEMBL, KEGG/GO) as well as specialized databases for Traditional Chinese Medicine (TCM)^14^, and use them to construct knowledge graphs that represent compound-gene-disease/phenotype relationships. These graphs then serve as the foundation for inferring novel links that conventional filters often overlook. This approach is consistent with recent reports highlighting the transformative role of AI and knowledge graphs in drug discovery in general^15^, and the growing application of artificial intelligence in natural product research specifically^16^.

Accordingly, we developed an early-stage modular, four-phase system that integrates biomedical data, knowledge graphs, and LLMs. It supports dual inference routes: forward (compound → gene → disease/phenotype) and reverse (disease/phenotype → gene → compound), and is designed for modular reuse across both directions. Integrating LLMs with knowledge graphs is emerging as a promising strategy to address inherent limitations of LLMs, such as factual unreliability and reasoning challenges, particularly in the demanding context of biomedicine^17^. A prime example is the recent development of Knowledge Graph-based Retrieval Augmented Generation (KG-RAG) frameworks, which leverage massive biomedical KGs to ensure that LLM-generated responses are rooted in established knowledge, thereby dramatically enhancing their reliability and accuracy for tasks like question answering^18^. Building upon this progress, our study explores a system designed to move beyond ranked lists or query responses, and instead generate explicit *natural language hypotheses*. The goal of this early experimentation is to enhance interpretability and support testable biological reasoning, thereby demonstrating the feasibility of this approach for complex conditions.

As a case study, we applied the system to Shakuyaku-kanzo-to (also known as Shaoyao-Gancao Tang in TCM)^19^. This formula was chosen because it consists of only two herbal components and is widely used in clinical practice with substantial supporting evidence. Indeed, Shakuyaku-kanzo-to has robust clinical documentation for its application in managing complex, inflammation-related pain syndromes associated with modern cancer therapies^20,21^. Using the early implementation of Phases 1 and 2, the system generated a hypothesis linking paeoniflorin to the immune receptor *TLR4*, which was not identified by conventional filtering.

Overall, this study demonstrates the system’s potential to process naturally derived multicomponent formulas such as Kampo, and underscores the feasibility of integrating LLMs and knowledge graphs to generate biologically plausible hypotheses for complex conditions. Unlike prior frameworks that mainly provide ranked targets or pathway lists, the present approach offers an important conceptual advance: the automated generation of structured, testable hypotheses articulated in natural language. This capability substantially improves interpretability and provides more direct support for biological reasoning.

### Notation

Throughout this paper, all gene and protein names are consistently italicized for clarity, regardless of conventional typographic distinctions.

## 2 Background and Related Work

### 2.1 Data-Driven Filtering Approaches

In network pharmacology, various data-driven filtering techniques, such as database-driven compound screening and *in silico* target prediction, have been widely adopted to reduce large-scale molecular datasets and facilitate the construction of interpretable interaction networks^22^. These approaches are widely adopted due to their computational efficiency and alignment with structured biomedical resources.

However, these techniques are fundamentally limited by the quality and completeness of the underlying data^23^. This creates a critical trade-off: strict thresholds systematically exclude compounds with limited annotation, such as paeoniflorin from *Paeoniae Radix*, while looser thresholds yield excessively large gene sets that are impractical for analysis. This bottleneck, where filtering either omits relevant entities or generates overwhelming complexity, highlights the need for novel approaches that can navigate sparse data^24–26^.

In this study, filtering is retained only as a minimal preprocessing step for noise reduction. The core analytical process instead combines structured knowledge graph integration with inference supported by LLMs. By leveraging textual associations and semantic reasoning, LLMs enable hypothesis generation even in the absence of explicit compound-target links, for example by extrapolating indirect associations between compounds, genes, and diseases or phenotypes. Thus, the pipeline shifts from threshold-based exclusion to structure-aware and language-informed inference, expanding the discovery space while preserving interpretability.

### 2.2 Network Pharmacology and Traditional and Naturally Derived Medicines

Recent advances in network pharmacology have enabled systematic analyses of the complex relationships among drugs, targets, and diseases. By integrating multiple resources, such as specialized databases for traditional medicine (e.g., TCMSP^14,27^) and general-purpose resources like STITCH and ChEMBL, researchers can model how compounds interact with proteins and signaling pathways at a systems level. This integrative approach has been particularly valuable for characterizing the multicomponent and multi-target properties of traditional and naturally derived medicines. It has also helped establish a powerful paradigm to understand their holistic effects^28,29^.

However, a significant portion of the published literature remains primarily descriptive. Many studies rely heavily on *in silico* predictions and culminate in extensive lists of potential genes or pathways. These studies often stop short of translating their complex association maps into testable, mechanistic hypotheses. This gap between computational prediction and experimental validation is a widely recognized challenge. Recent reviews emphasize that the results of *in silico* analyses must be validated by experimental findings to be meaningful^30^. This underscores the need for frameworks that can move beyond descriptive mapping. Such frameworks should bridge the gap from association to explanation by generating specific, testable hypotheses that can directly guide laboratory research.

### 2.3 AI-Driven Hypothesis Generation

Recent advances in artificial intelligence (AI) offer new opportunities to address the limitations of conventional data-driven filtering. Domain-specific language models such as BioBERT^31^ and SciBERT^32^, trained on biomedical and scientific corpora, can extract and integrate knowledge dispersed across literature and databases. More recently, the powerful ability of LLMs like the GPT family^33^ to synthesize diverse information offers the potential to articulate complex hypotheses in natural language. However, these AI-driven approaches face significant challenges, including unsupported associations (hallucination) and difficulties in validation^34,35^. Thus, their scientific utility depends on careful integration with structured knowledge bases to ground their reasoning. This study explores such an integrative approach, proposing a pipeline where the primary output is an explicit, natural-language hypothesis (e.g., Compound X may affect Disease Y via Pathway Z) designed to inform subsequent research.

### 2.4 Integrative Efforts: Toward Multimodal Medicine

Integrative efforts bridging traditional and naturally derived medicines with modern pharmacology have been reported, particularly under network pharmacology and systems pharmacology frameworks that analyze the multi-target mechanisms of herbal formulations^1,5^. Recent perspectives also emphasize the need for multimodal strategies that combine heterogeneous data sources to uncover complex biological mechanisms^34,36^. Building upon these efforts, the present study aims to fill the gap between data and explanation by proposing a system that leverages the integrative reasoning capabilities of LLMs to generate natural language hypotheses, thereby bridging natural compound–based and contemporary pharmacology.

## 3 System Vision and Architecture

We propose a modular system for hypothesis generation that integrates structured biomedical databases, knowledge graphs, and language models into four interconnected phases aimed at supporting drug discovery. As summarized in Figure 1, the system is organized into four interdependent phases, supported by biomedical databases and knowledge graphs^37^.

**Figure 1.**
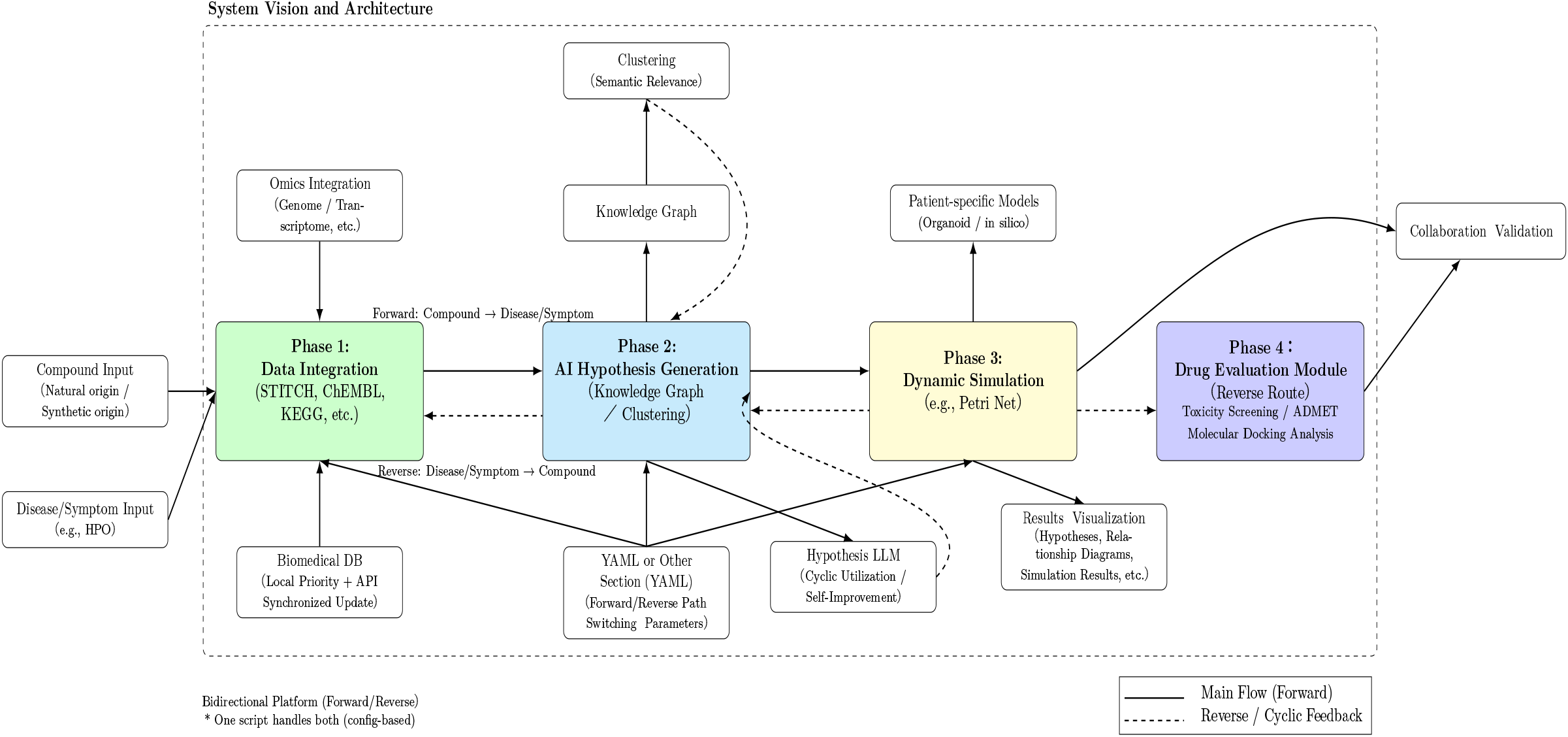
System Vision and Architecture of the Modular Four-Phase Hypothesis Generation Pipeline. Phase 1: Aggregates and scores biomedical data from multiple domains. Phase 2: Generates hypotheses via character sheets, knowledge graphs, clustering, and semantic/biological screening. Phase 3: Applies dynamic Petri net simulation to assess causal and temporal plausibility. Phase 4: Performs *in silico* drug discovery evaluation (docking, ADMET, and drug-likeness). The system supports both forward (compound → gene → disease/phenotype) and reverse (disease/phenotype → gene → compound) inference routes, enabling both novel discovery and repurposing. A modular, self-updating framework that supports integration of diverse compound modalities, spanning natural products and modern pharmaceuticals, including but not limited to traditional formulations such as Kampo.

### 3.1 Modular Four-Phase System Architecture

The system is designed to accommodate hypothesis generation in multi-target and multimodal pharmacological settings. While Kampo medicine served as an initial test case, the architecture is intentionally generalized to support various compound modalities, including polyherbal formulations, small molecules, and biologics. This flexibility enables use across both naturally derived and modern therapeutic pipelines.

- **Phase 1: Data-Driven Aggregation and Scoring** Open biomedical resources covering multiple domains, such as compound–target interactions, biological pathways, protein-protein interactions, non-coding RNA associations, tissue-specific expression, and disease-phenotype relationships, are integrated. Whenever possible, datasets are pre-downloaded, pre-processed, and stored locally; API access is used for resources without bulk downloads. A latent information score is computed per gene based on evidence density and source diversity, resulting in a unified per-gene information sheet for downstream use.
- **Phase 2: Hypothesis Generation via Semantic and Biological Reasoning** This phase transforms prioritized data from Phase 1 into structured and mechanistic hypotheses. It is organized into several core modules that progressively refine compound–gene–disease/phenotype associations into higher-order mechanistic insights:
  – **Character Sheet Generation:** For each high-priority gene, the system generates a semantic *Character Sheet (CS)* summarizing the gene’s biological roles, interactions, and disease or compound associations. GPT-based models are currently used, but the design already supports future multimodel consensus to improve robustness and minimize bias.
  – **Knowledge Graph Construction and Clustering:** Following established methodologies^37^, genes, compounds, and diseases are integrated into a knowledge graph representing known and inferred relationships. The graph is partitioned using the Leiden algorithm to identify modules that may reflect biological pathways or therapeutic mechanisms^38^.
  – **Cluster-Level Hypothesis Extraction:** Within each cluster, gene-level CS are aggregated and submitted to LLM agents for synthesis. Prompts encourage mechanistic reasoning about disease processes and therapeutic effects.
  – **Formula-Level Hypothesis Synthesis:** Cluster hypotheses can be further integrated into higher-level *formulas*, corresponding to compound mixtures or empirical formulations. This allows flexible connections between molecular mechanisms and treatment paradigms. Modules such as semantic screening with BioBERT^31^, SciBERT^32^, and PubMedBERT^39^ remain under development, and will be systematically benchmarked in future iterations. Future work will extend this phase toward more robust hypothesis selection and evaluation.
- **Phase 3: Dynamic Simulation with Petri Nets** Petri nets are used to simulate and evaluate the temporal and causal plausibility of hypotheses, a method widely applied in systems biology^40,41^. Beyond pathway-level applications, Petri net modeling has also been extended to integrative representations in East Asian medicine, such as acupuncture and moxibustion^42^. Matsuno and colleagues have contributed theoretical foundations of Petri nets in systems biology^43^, while Ge and collaborators have explored their application to traditional medicine contexts. Together, these complementary efforts highlight the versatility of Petri nets in capturing complex biological processes, making them a suitable option for future implementation of Phase 3. Top-ranked hypotheses (3–5) are prioritized for simulation to balance feasibility and interpretability. This provides:
  – Testing of hypothesized mechanisms under dynamic conditions,
  – Identification of emergent behaviors such as activation, inhibition, or compensatory feedback, and
  – Highlighting of critical nodes or transitions as potential intervention points. This phase bridges static knowledge aggregation and dynamic biological reasoning.
- **Phase 4: Drug Discovery Evaluation** Although not yet implemented, this phase will provide a lightweight *in silico* evaluation pipeline including:
  – **Molecular Docking:** Assessing compound binding compatibility with target proteins^44^.
  – **ADMET Profiling:** Estimating absorption, distribution, metabolism, excretion, and toxicity^45^.
  – **Drug-Likeness Filtering:** Applying cheminformatics rules (e.g., Lipinski’s Rule of Five^46^). This stage acts as an early filter to prioritize promising candidates rather than replicate full-scale drug development.

### 3.2 Dual Inference Routes and Modular Reusability

The system supports two complementary inference pathways:

- **Compound-driven Route** (Compound → Gene → Disease/Phenotype): Starting from known compounds to uncover new indications.
- **Disease-driven Route** (Disease/Phenotype → Gene → Compound): Starting from a disease or phenotype to suggest candidate compounds.

Both routes share a unified modular architecture. By interchanging YAML-based configuration templates, core modules (e.g., knowledge graph, clustering, hypothesis extraction) can be reused seamlessly.

### 3.3 Key Design Features

Key architectural features include:

- **Self-Updating Knowledge Base:** Local caching and API retrieval for reproducibility and cumulative expansion.
- **Graph Reasoning and Unsupervised Clustering:** Leiden clustering is prototyped; alternative algorithms will be benchmarked to improve robustness.
- **Support for Integrative Medicine:** The system is adaptable to patient-specific models and may map Kampo diagnostic categories to biomedical ontologies in the future.
- **LLM-Based Hypothesis Memory:** High-confidence hypotheses could be archived to fine-tune domain-specific LLMs, creating a feedback loop for refinement.

These phases together establish a reproducible workflow that progresses from raw biomedical data to mechanistic hypotheses and preliminary validation. Ultimately, the modular pipeline forms a bridge from raw biomedical data to hypothesis-driven drug discovery, supporting both rational design and exploratory research.

## 4 Implementation

This section outlines the current implementation status based on the modular, four-phase architecture introduced in the **System Vision and Architecture**. The present study primarily covers Phase 1 and parts of Phase 2, providing an early-stage foundation for hypothesis generation via knowledge graph analysis. As such, this work represents an initial implementation toward a scalable, extensible system.

### 4.1 Phase 1: Data-Driven Knowledge Construction

In Phase 1, compounds were first retrieved from KampoDB^47^, a Japanese database that catalogs traditional formulas and crude drugs together with associated information such as constituent compounds, target proteins, and pathway annotations (e.g., KEGG^48,49^ and GO^50^). Based on the herbal components of each formula, compound entries from KampoDB were collected and then linked to target proteins mainly through STITCH^51^ and ChEMBL^52^, which provide complementary coverage: STITCH for bulk historical data and ChEMBL for recent, structured annotations. Additional biomedical databases covering pathways, protein-protein interactions, gene expression, and disease-phenotype relationships were also incorporated as available (see also Figure 2A and Supplementary S1).

**Figure 2.**
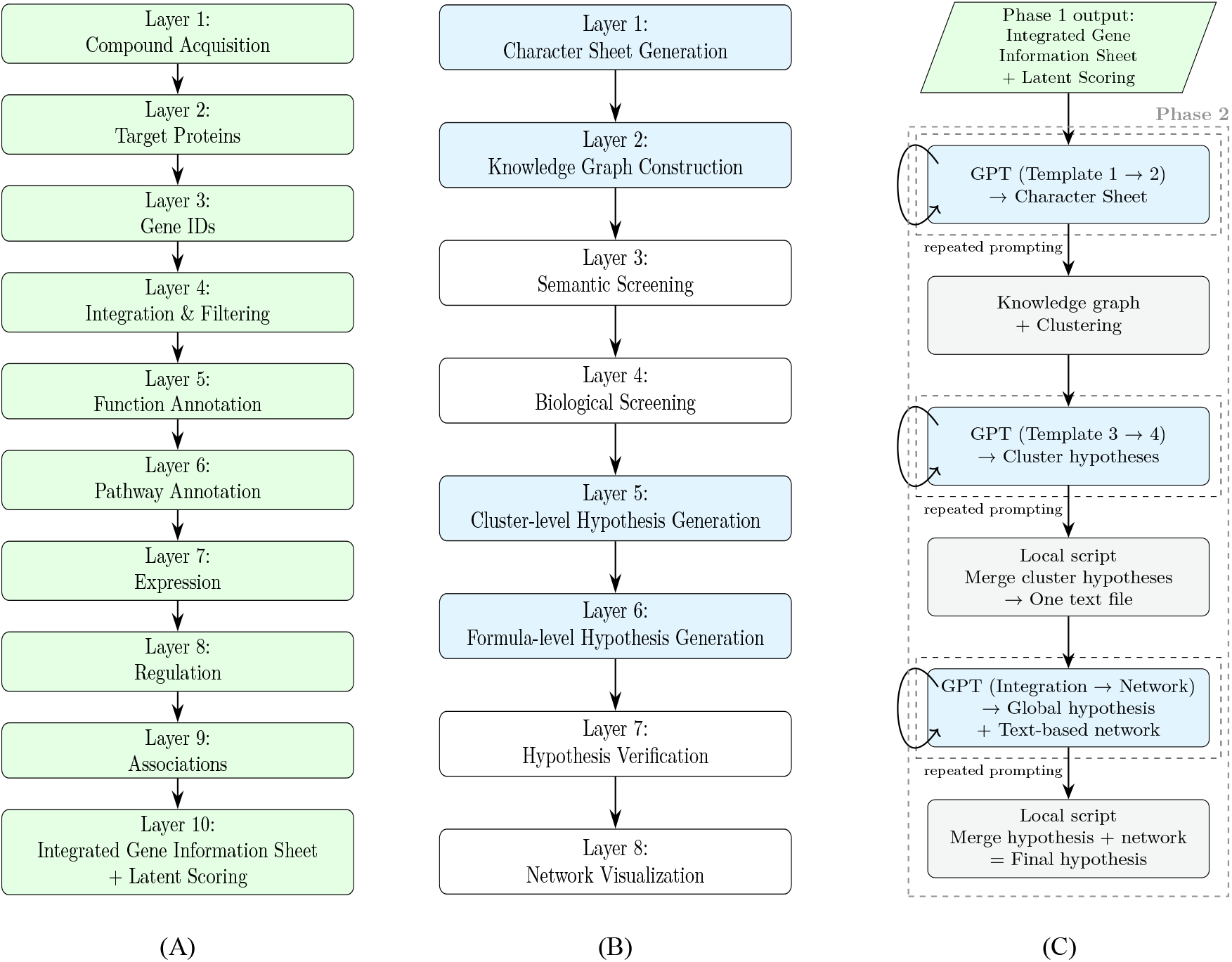
Forward information flow across two phases. Each layer represents a logical step in the pipeline. (A) Phase 1: Data Integration Layered Structure Overview (B) Phase 2: Hypothesis Generation Layered Structure Overview (C) Phase 2: Implemented forward route (initial implementation only) *Note: Layer shading indicates early-stage implementation. Full implementation details of Phase 1 and Phase 2 are summarized in Supplementary Tables S1 and S2, respectively*.

For data sources without bulk downloads, such as certain disease ontologies, API-based access was implemented. To ensure both efficiency and reproducibility, a self-updating, self-growing local database was developed: each query first checks the local repository, and only if data are missing, the system retrieves and stores them externally. Over time, this approach builds a cumulative knowledge base while minimizing redundant queries.

Each gene is associated with a structured Gene Information Sheet (GIS) that integrates multiple evidence types (e.g., compound-gene links, pathways, disease/phenotype associations). A latent score is assigned to each gene based on the quantity and diversity of evidence, enabling prioritization while preserving transparency.

### 4.2 Phase 2: Hypothesis Generation (Partial Implementation)

Phase 2 focuses on the initial generation of hypotheses from the integrated knowledge base. Some modules have been implemented, while others remain under development (see also Figure 2B and Supplementary S2).

- **Character Sheet Generation (Implemented, Prototype):** A semantic Character Sheet (CS) is created for each prioritized gene, currently implemented using GPT-based models. This prototype stage was designed to first test whether informative outputs can be generated from a single model. Once feasibility was confirmed, the module was structured to allow extension into a multimodel consensus framework (e.g., Gemini, Claude), which will help improve robustness and minimize model-specific bias in future iterations.
- **Knowledge Graph Construction and Clustering (Partially Implemented):** CSs are converted into triples (subject-relation-object) and organized into a knowledge graph, where genes, compounds, and diseases are represented as interconnected entities. The resulting graph is partitioned using the Leiden community detection algorithm^38^ to identify cohesive modules that form the basis for downstream hypothesis generation.
- **Cluster-Level Hypothesis Generation (Implemented, Early):** As depicted in Figure 2C, Phase 2 involves four sequential LLM calls. First, GIS are abstracted into CS through a two-step prompting process: the initial LLM output is refined via a template-guided second call. Second, cluster-level hypotheses are synthesized in a similar two-step manner: a subset of CSs (top N genes by cluster subgraph centrality) is concatenated, submitted to the LLM, and refined through an additional template-based call. This design adopts an iterative prompting approach to improve coherence and interpretability. Prompts are structured to encourage mechanistic reasoning about how the selected gene set may contribute to disease processes or therapeutic effects.
- **Formula-Level Hypothesis Generation (Implemented, Early):** Multiple cluster-level hypotheses can be combined into broader formula-level hypotheses, enabling cross-cluster reasoning and higher-level integration.

Although hypothesis generation is already functional, additional modules are under development to enhance precision and reliability. These include semantic screening (planned using BioBERT, SciBERT, and PubMedBERT) and functional pathway layering prior to generation, as well as hypothesis verification after generation. Future extensions also include visualization and simulation modules to further improve interpretability.

### 4.3 Prompting Strategy and Template Design

In this implementation, six distinct prompting templates were designed and applied in a staged manner to structure the interaction with LLMs. Two templates were used for CS generation, supporting stepwise abstraction from the GIS. Two templates were employed for cluster-level hypothesis generation, enabling iterative refinement of hypotheses within each Leiden-derived cluster. Finally, two templates were applied for the integrative synthesis step, combining the 35 cluster-level outputs into both (i) a unified textual summary and (ii) a network-style structural representation.

This modular prompting strategy reflects a practical approach for managing complexity and guiding LLM-based reasoning in hypothesis generation. While template details are implementation-specific, their staged use illustrates how generative models can be integrated into a reproducible pipeline.

### 4.4 Case Study: Shakuyaku-Kanzo-to

As a case study, we used the Kampo formula *Shakuyaku-kanzo-to*^19^, selected for its simplicity (two herbs) and strong clinical evidence in anti-inflammatory applications.

Even such a small prescription yielded 220 unique compounds (3 from *Shakuyaku* and 217 from *Kanzo*), a result that highlights the intrinsic multicomponent nature of Kampo. From these, 68 compounds (2 from *Shakuyaku* and 66 from *Kanzo*) remained after integration with STITCH and ChEMBL, which were then mapped to protein targets. Integration with UniProt^53^, GO, KEGG, Reactome^54^, and other resources subsequently yielded 913 unique genes. For each gene, a GIS was generated, and a latent score was calculated based on evidence density and diversity. Among these, 123 genes scored ≥ 70 and were prioritized for Phase 2 processing.

In Phase 2, these high-priority genes were used for knowledge graph construction, clustering, and hypothesis generation. Detailed results, including cluster statistics and selected hypotheses, are presented in the **Preliminary Results** section.

To support the development and documentation of this system, we used standard programming tools and language editing assistants for code management and manuscript refinement.

## 5 Preliminary Results

### 5.1 Effects of Multi-Stage Prompting

To support hypothesis generation, we employed a multi-stage prompting strategy involving abstraction, synthesis, and network-based reasoning.

Initially, gene-specific knowledge was collected into comprehensive GIS (one per gene) from the Phase 1 data-driven layer. Through a two-pass GPT refinement process, these sheets were transformed into structured CS that summarized each gene’s context and functional role.

Based on these CS, a gene interaction network was constructed and subjected to Leiden clustering. In this case study, resolution parameters produced 35 clusters, each subsequently processed with a standardized prompt template in the compound, gene, and disease directions, designed to elicit plausible mechanistic hypotheses. This procedure was applied uniformly across all clusters.

The 35 cluster-level hypotheses were then synthesized into a unified *meta-hypothesis*, hereafter referred to as such, using two additional GPT steps. The first step integrated the textual content of individual hypotheses, while the second generated a graph-style structural summary to capture inter-gene relationships. The resulting composite output combined textual synthesis with network visualization and was treated as the final hypothesis for downstream analysis in this study.

### 5.2 Cluster Statistics

We applied the Leiden clustering algorithm to the structured gene interaction network derived from 123 high-scoring genes (latent score ≥ 70). The resolution parameter was varied from 0.3 to 14.0, with higher values yielding progressively more clusters (Figure 3).

**Figure 3.**
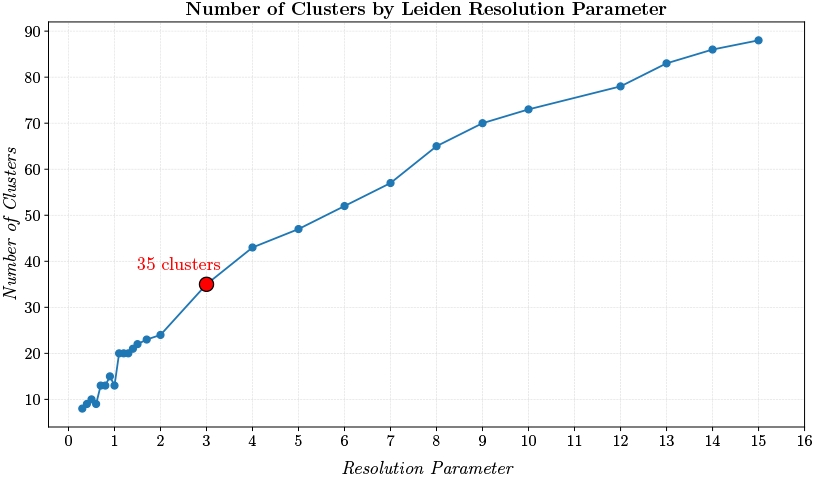
Cluster Count vs. Resolution: Number of clusters generated by the Leiden algorithm across different resolution parameters. At resolution 3.0 (35 clusters), *TLR4* emerged in a distinct cluster. At higher resolutions, it was occasionally reassigned to other upstream groupings, reflecting parameter sensitivity.

At a resolution of 3.0, the algorithm produced 35 clusters, a practical balance given the parameter sensitivity of Leiden clustering^38^. For example, it was the first point at which the gene *TLR4*, a key upstream receptor in innate immunity^55^, appeared in a distinct cluster. At lower resolutions, *TLR4* was absorbed into larger modules, whereas higher resolutions fragmented the network into many small clusters. Beyond 3.0, *TLR4* did not consistently remain isolated, reflecting the sensitivity of Leiden clustering to parameter choices and local network structure. Thus, resolution 3.0 is used here as a practical working point for hypothesis generation, rather than as an absolute optimal value.

### 5.3 Hypothesis Emergence

Among the resulting clusters, Cluster 14 was highlighted as a representative case. This cluster contained five key player genes with explicit ranks: *NFKBIA* (rank 0, highest), *TLR4* (rank 1), *NFKB1* (rank 2), *CHUK* (rank 3), and *RELA* (rank 4). Although *TLR4* was not the top-ranked node, it was highlighted because paeoniflorin, a compound often excluded by conventional filtering, was identified as being linked to this receptor through hypothesis generation. In the present framework, both the ranked genes and their associated compounds/diseases are treated as *key players* within each cluster.

At the chosen resolution of 3.0 (35 clusters), hypotheses were generated using GPT-based agents. A general-purpose forward-route prompt template, designed to explore compound-gene-disease relationships, was uniformly applied to all clusters. Each cluster thus produced a structured hypothesis, and we subsequently focused on the hypothesis from Cluster 14 due to the presence of *TLR4*.

The generated hypothesis from Cluster 14 included the following representative sentence:

> “Paeoniflorin and Puerarin are hypothesized to modulate *TLR4* activity, potentially attenuating inflammatory responses associated with depression and atherosclerosis by dampening NF-κB and HIF-1 signaling”.

This aligns with existing literature describing paeoniflorin’s anti-inflammatory effects^56^, puerarin’s pharmacological activities^57^, and the established roles of NF-κB and HIF-1 in inflammation^58,59^. The compound-gene–disease interaction structure for Cluster 14 is shown in Figure 4.

**Figure 4.**
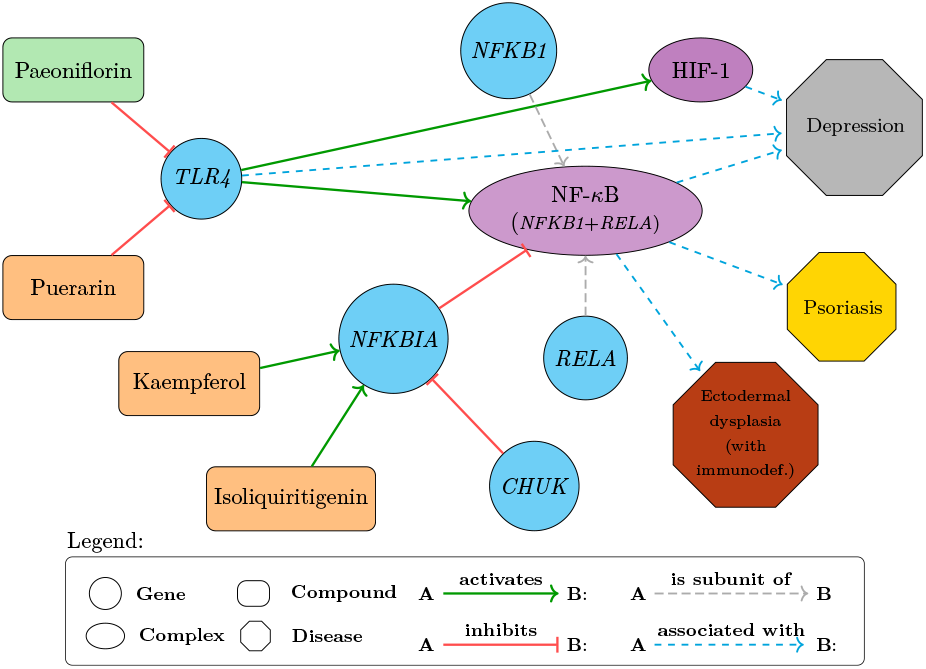
Cluster 14 Inference Network: This network was constructed according to the structured hypothesis generated for Cluster 14, with *NFKBIA* and *TLR4* as central genes. All depicted relations were manually verified against PubMed, as documented in Supplementary Table S5. Nodes represent genes, compounds, or diseases as indicated in the legend.

Although *TLR4* was not the top-ranked node in Cluster 14, its role as an upstream receptor within the NF-κB axis, and its specific emergence through hypothesis generation, warranted further attention. A full version of the hypothesis is provided in Supplementary S3.

### 5.4 Final Hypothesis Integration

This final hypothesis was developed as an integrative summary of the 35 cluster-level outputs, aiming to provide a broader systems-level perspective on how the components of the herbal formula *Shakuyaku-kanzo-to* may exert biological effects across multiple pathways.

Rather than focusing on individual clusters, this hypothesis organizes the findings into four conceptual modules: Inflammation & Immune Modulation, Neuro-Psychiatric Modulation, Metabolic & Endocrine Modulation, and Anti-Tumor & Cell Proliferation. The grouping was based on shared compounds, pathway overlaps, and recurring gene targets identified during analysis.

Importantly, the exemplars in this integrated hypothesis are grounded in the cluster-level outputs rather than post hoc curation. For instance, genistein acting on the *IL6*–*TNFRSF1A* axis (Cluster 17), glycyrrhiza acting on the *CXCR4*–*MIF* axis (Cluster 13), genistein modulating *CYP3A4* (Cluster 31), glycyrrhiza and genistein engaging *PTH1R* (Cluster 33), genistein targeting *RET* (Cluster 34), melatonin modulating *MTOR* (Cluster 27), and serotonin/glycyrrhiza and daidzein acting via *SLC6A4*/*HTR1A* and *GABRG2* (Cluster 23) are explicitly reflected in the module summaries (see Supplementary S4 for full text).

Key regulatory hubs such as *IL6, MTOR*, and *CYP3A4* appeared across multiple modules, suggesting that they may play central roles in mediating broad-spectrum effects. Likewise, compounds like genistein, glycyrrhiza-derived constituents, and melatonin were observed in multiple functional contexts, indicating potential multifunctionality.

An emphasis is placed on network crosstalk, where compounds may act across modules via shared targets. For example, genistein is involved in inflammatory, metabolic, and anti-tumor processes, while glycyrrhiza contributes to immune, psychiatric, and endocrine-related effects.

This summary complements the cluster-level hypotheses by providing a high-level map situating them within a broader framework. It may be viewed as a synthetic perspective that translates a traditional Kampo formula into a modern molecular narrative.

### 5.5 Evaluation and Validation

The primary objective of this study is to present the overall system design and its initial implementation; a systematic validation of the generated hypotheses has not yet been undertaken. Nevertheless, validation is essential, and our current idea can be summarized as follows:

i. Extract triples in the form of (subject-relation-object), such as (compound-activates-gene) or (gene-associated_with-disease), from hypotheses written in natural language,
ii. Search for supporting evidence in biomedical literature databases (e.g., PubMed),
iii. Interpret the outcomes: if evidence is found, the hypothesis is considered plausible; if not, the case may represent either *novelty* or *error*.

**Table 1.**
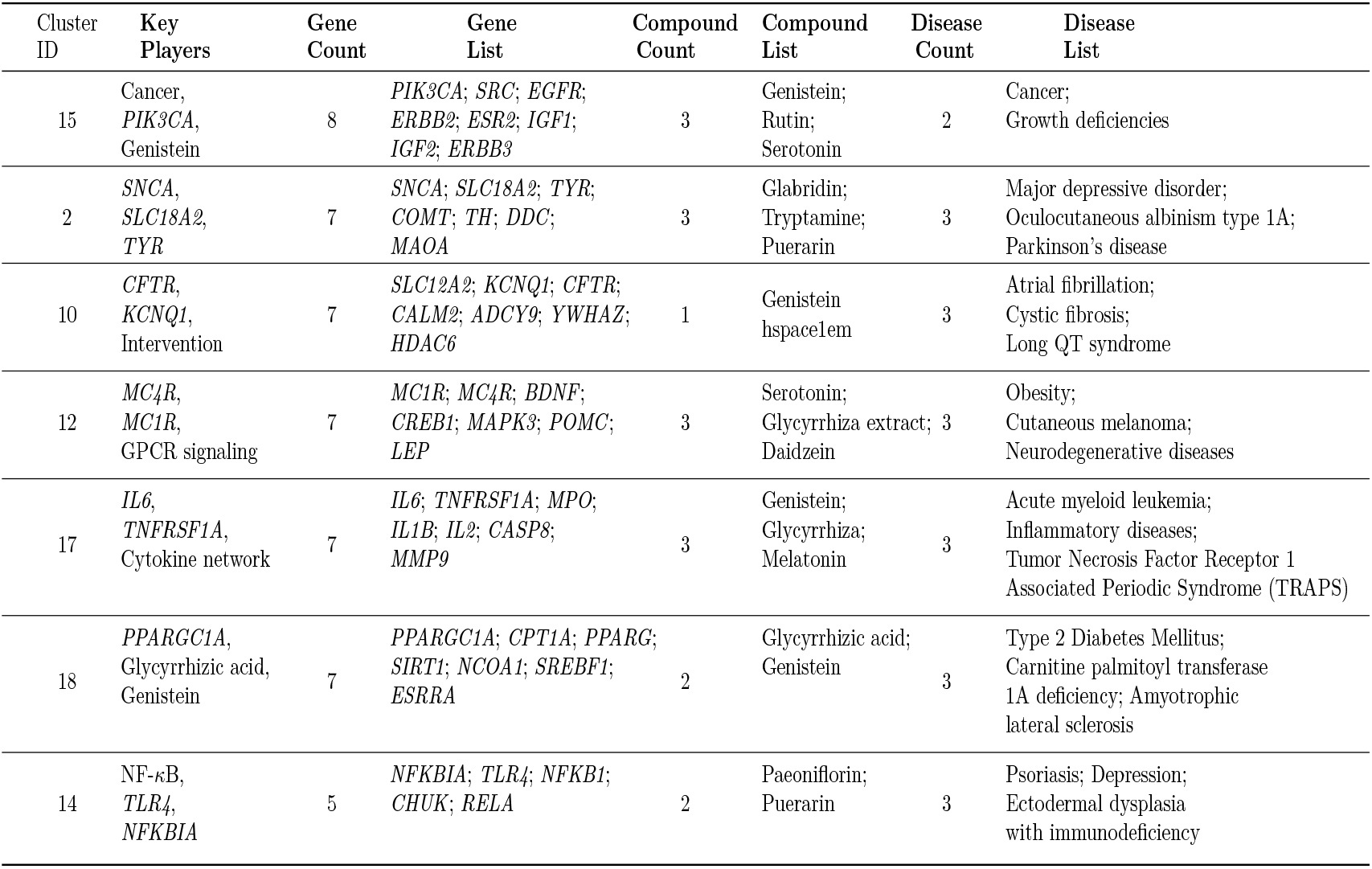
Composition Summary for Major Clusters. Clusters are ordered by gene count (≥ 7). Cluster 14, despite its moderate size, is included due to its biological relevance, particularly involving *TLR4*. For Cluster 14, all depicted relationships were manually cross-checked against PubMed literature.

As an illustrative case, we manually validated Cluster 14. All triples shown in Figure 4 were checked against PubMed, and no contradictions with existing literature were identified. These findings suggest that the generated hypotheses were not arbitrary and, at least in this cluster, align with established biological knowledge.

Importantly, the hypotheses examined here were based on the top 123 genes (latent score ≥ 70) out of 913 identified in Phase 1. Looking ahead, in Phase 2 (Layer 7) we plan to formalize this validation framework and implement automated pipelines that can systematically assess plausibility, novelty, and error.

## 6 Discussion

### 6.1 System-Level Capabilities and Limitations

- **Strengths:** The system identified coordinated gene relationships, such as the co-modulation of *TLR4* and *NFKBIA*, both key regulators in inflammatory signaling. Several extracted compounds, including paeoniflorin and puerarin, aligned with known Kampo formulations, lending further plausibility to the generated hypotheses. Certain hypotheses extended beyond single-target interactions, suggesting that the system may capture multi-layered and system-wide biological regulation.
- **Limitations and Challenges:** Several limitations remain. The current implementation lacks temporal reasoning and confidence scoring, a common issue in early systems pharmacology pipelines^60^.

LLM-specific issues were also observed. Token constraints occasionally truncated outputs, and repeated prompting introduced semantic drift. In some cases, the system produced non-existent entities, highlighting the need for robust entity normalization.

Prioritization inconsistencies were noted as well. In Cluster 14, domain experts identified *NFKBIA* as central, yet the model emphasized *HIF1A*, whose relevance was less clear. These discrepancies underscore the importance of human-in-the-loop refinement for biologically grounded interpretations.

Finally, while the system generated plausible compound-gene-disease associations, it has not yet provided detailed mechanistic explanations or temporal dynamics. Addressing these aspects will be critical for advancing hypothesis generation into actionable scientific insight.

### 6.2 Technical Design Insights: Clustering and Prompt Engineering

- **Clustering Approaches:** Initial implementations relied on Louvain clustering, but the results were inconsistent and heavily affected by initialization randomness. Leiden clustering was adopted as a replacement, offering more stable modular divisions and improving the interpretability of gene communities. Although this transition did not directly yield key hypotheses (e.g., the *TLR4*-NF-κB axis), it improved the overall stability of modules across parameter settings. Further refinements are needed. For example, enhancing the knowledge base by moving from triple-based graphs to richer supergraph representations could improve clustering coherence. Clustering outputs may still be affected by biases or artifacts in the input data. Therefore, benchmarking alternative algorithms and refining graph construction, such as incorporating multi-relational edges or transitioning toward supergraph representations, remains an ongoing priority.
- **Prompt Design Considerations:** Prompt engineering played a critical role in shaping outputs. For example, when templates required the model to enumerate a fixed number of items (e.g., two compounds or three diseases), the model occasionally skipped required elements, producing incomplete results. We refined the templates with stricter constraints, which reduced such obvious errors. Nevertheless, hidden errors may still remain, underscoring the need for systematic validation mechanisms beyond prompt design. While natural language hypotheses offer the advantage of presenting fragmented biomedical knowledge in a more connected and interpretable form, they also introduce ambiguity compared to triple-based representations. We therefore regard natural language not as a replacement, but as a complementary layer: a way to generate integrative hypotheses that can later be validated and refined through structured knowledge and experimental evidence.

### 6.3 Reconsidering the *Jun*-*Chen*-*Zuo*-*Shi* Model in the Context of Shakuyaku-kanzo-to: A Hypothesis Centered on TLR4

Based on the AI-generated hypothesis, we constructed and visualized a compound-gene-disease interaction network (Figure 4) to assess the emergence of interpretable pharmacological patterns. The resulting network suggested that paeoniflorin from *Paeoniae radix* targets the upstream receptor *TLR4*, while constituents from *Glycyrrhizae radix* (e.g., kaempferol and isonaringin) act on downstream effectors such as *NFKBIA*. This upstream-downstream arrangement echoes the traditional *Jun*-*Chen*-*Zuo*-*Shi* (君 臣佐使, JCZS) framework in Kampo medicine, which originates from TCM^6,19^. In this framework, the *Jun* (君) addresses the core pathology, while the *Chen* (臣) supports and modulates the therapeutic effect.

However, the inclusion of puerarin complicates this interpretation. According to KampoDB, puerarin, assigned to *Glycyrrhizae radix* (Kanzo), was likewise predicted to target the same upstream receptor *TLR4* as paeoniflorin. Subsequent validation confirmed that puerarin does indeed modulate *TLR4*. While both paeoniflorin and puerarin were predicted to act on the upstream receptor *TLR4*, the assignment of puerarin to *Glycyrrhizae radix* remains to be further substantiated.

In the classical framework, the *Jun* is traditionally regarded as the principal agent that suppresses the main pathology, and the *Chen* plays a complementary role. In this case, simultaneous targeting of *TLR4* by both paeoniflorin and puerarin blurs the distinct roles envisioned in the JCZS framework, suggesting that the classical notion of a singular *Jun* may not apply cleanly when multiple constituents act on the same upstream node.

We do not regard this as a contradiction of the JCZS framework. Rather, it calls for a reinterpretation of traditional roles in light of emerging molecular evidence. This case illustrates a mismatch between the classical JCZS framework and the observed molecular upstream-downstream relationships. However, this observation alone is insufficient to reject either the molecular pathway model or the JCZS paradigm. Further investigation, including dynamic simulation and broader case studies, is needed to evaluate their possible alignment. Importantly, while Kampo provides a clear illustration, the same framework can be applied to other natural compound–based therapies beyond traditional contexts.

### 6.4 Toward Robust Validation

While this study was able to generate multi-layered hypotheses using language models and clustering techniques, rigorous validation remains a critical next step. At present, validation was limited to manual PubMed queries that confirmed individual relationships (e.g., paeoniflorin-*TLR4, NFKBIA*-inflammation), but did not extend to integrated system-level hypotheses. This limitation underscores the importance of implementing structured, scalable validation mechanisms in the next phase.

In particular, several components of the Phase 2 architecture remain under development:

i. **Layer 3: Semantic Screening** will incorporate models like BioBERT or SciBERT to refine the biological plausibility of intermediate entities and relationships.
ii. **Layer 4: Functional Layering** aims to align hypotheses with physiological pathways and logical frameworks.
iii. **Layer 7: Hypothesis Validation** is critical for determining the accuracy and usability of generated hypotheses through evidence-based reasoning.

These layers are prioritized for upcoming implementation before moving toward any dynamic simulation (i.e., Phase 3). Validation is not an auxiliary task but an essential structural component to ensure the utility of hypotheses generated in Phase 2.

Furthermore, the quality of validation depends heavily on the strength of the underlying knowledge graph and clustering mechanisms (Layer 2). Although partially implemented, these components require further refinement. Specifically:

- A shift toward supergraphs (richer graph structures with multiple entity types and annotated edge semantics) is being considered to represent richer, multi-relational biomedical contexts.
- Clustering methods must be evaluated and potentially replaced or improved to ensure robust and biologically meaningful module generation.

Taken together, these planned enhancements will enable both quantitative and qualitative validation of hypotheses and provide a reliable bridge toward Phase 3 simulations.

### 6.5 Future Directions and Integration

Several extensions are envisioned to enhance the quality of hypotheses and improve system robustness:

- **Multimodel consensus:** In Phase 2, multiple language models (e.g., GPT, Gemini, Claude) will be incorporated to encourage convergence and reduce model-specific bias. Semantic screening layers (e.g., BioBERT) are also planned in parallel and could be integrated using ensemble methods.
- **Self-evolving framework:** By archiving hypotheses, prompts, and validation feedback, a dedicated training corpus could be developed for a specialized *Hypothesis LLM*. This feedback loop is expected to gradually transition the system from a generative assistant to a semi-autonomous scientific collaborator.
- **Integration with medicine and traditional knowledge:** The framework may be linked with personalized medicine, for instance by connecting patient-derived iPS data to gene-compound networks for individualized prediction. Similarly, Kampo diagnostic patterns (*Sho*) could be mapped to phenotype ontologies^1,5^ such as HPO, facilitating translation between empirical prescriptions and molecular mechanisms.
- **Toward multimodal drug discovery:** While the present study used a Kampo formula as a prototype, the same framework could be applied to future multimodal drug designs that combine multiple natural compounds with a small number of Western drugs. In such cases, the cluster-level hypotheses may more directly converge into an integrated final hypothesis, reflecting practical settings where fewer compounds are involved but synergistic interactions are critical. This direction highlights the potential of the framework as a bridge between natural product–based modalities, including but not limited to traditional medicines, and modern pharmaceuticals within a multimodal drug discovery paradigm.

Together, these directions aim to advance the system toward a scalable and semantically grounded platform for hypothesis-driven biomedical discovery.

## 7 Conclusion

This study introduces a modular, four-phase hypothesis generation system that integrates structured biomedical data, knowledge graphs, and language models to support early-stage drug discovery. As a case study, we applied the system to the Kampo formula Shakuyaku-kanzo-to and generated explicit natural language hypotheses, including associations such as paeoniflorin with the immune receptor *TLR4*, which were overlooked by conventional filtering approaches.

Cluster 14 was selected for preliminary validation, where all subject-relation-object triples (e.g., paeoniflorin-inhibits-*TLR4*) were manually checked against PubMed. No contradictions with the literature were found, suggesting biological plausibility and consistency with current knowledge. Although this constitutes only an early validation, it demonstrates the feasibility of natural language hypothesis generation as a complement to database-driven filtering.

A distinctive feature of the system is its support for dual inference routes (compound → gene → disease/phenotype and disease/phenotype → gene → compound), which enables both forward mechanistic exploration and reverse drug repurposing. This dual perspective, combined with language-informed reasoning, expands discovery beyond association mapping toward interpretable and testable biological hypotheses.

Looking ahead, the next priority is to implement systematic validation frameworks for language model–generated hypotheses, followed by dynamic Petri net simulation (Phase 3) and *in silico* drug evaluation (Phase 4). These steps will establish a self-refining pipeline, advancing from hypothesis generation to predictive modeling and candidate prioritization. If traditional multicomponent formulas such as Kampo can be systematically analyzed, this framework can extend into true *multimodal drug discovery* that integrates Western pharmaceuticals with naturally derived therapeutics.

In summary, this work presents an early proof of concept in AI-driven hypothesis generation. By introducing explicit, interpretable natural language hypotheses supported by dual inference routes, the system provides a foundation for future strategies that bridge natural product–based therapeutics and conventional one-drug-one-target pharmaceuticals within a unified multi-modal paradigm. Such an exploration might have been impractical in the past, but with the emerging reasoning capabilities of modern LLMs, it may now be possible to link fragmented knowledge into coherent hypotheses, suggesting new avenues for multimodal drug discovery.

## Data availability

The raw data used in this study are summarized in Supplementary S1. The processed datasets are available in Zenodo repository at https://doi.org/10.5281/zenodo.17043217. These include cross_cluster_genes.csv (35 clusters), cluster_hypotheses.zip (containing 35 JSON files named cluster_0_summary.json to cluster_34_summary.json), and final_hypothesis_report.md.

## Author contributions statement

R.W. led the conceptual design of the system, carried out all software implementation, and authored the manuscript. H.M. contributed biological expertise and supported the hypothesis validation. Q.G. contributed to the development and refinement of the system architecture. All authors reviewed the manuscript.

## Supplementary Materials

### S1 Phase 1: Data-Driven Knowledge Construction (Implementation Details)

This section provides a detailed list of databases and resources used in Phase 1 of the system implementation. For each resource, we indicate whether it is currently implemented or planned for future integration, along with the date of access and version number when available. These supplementary tables are intended to ensure transparency and reproducibility of the system, complementing the concise description in the main text.

#### Stage Definitions for Data Integration

- **Stage 1**: Fully implemented and used in the current system.
- **Stage 2**: Partially implemented or limited use.
- **Stage 3**: Data accessed and prepared, but not used in this implementation.
- **Stage 4**: Planned for future integration; data not yet accessed.

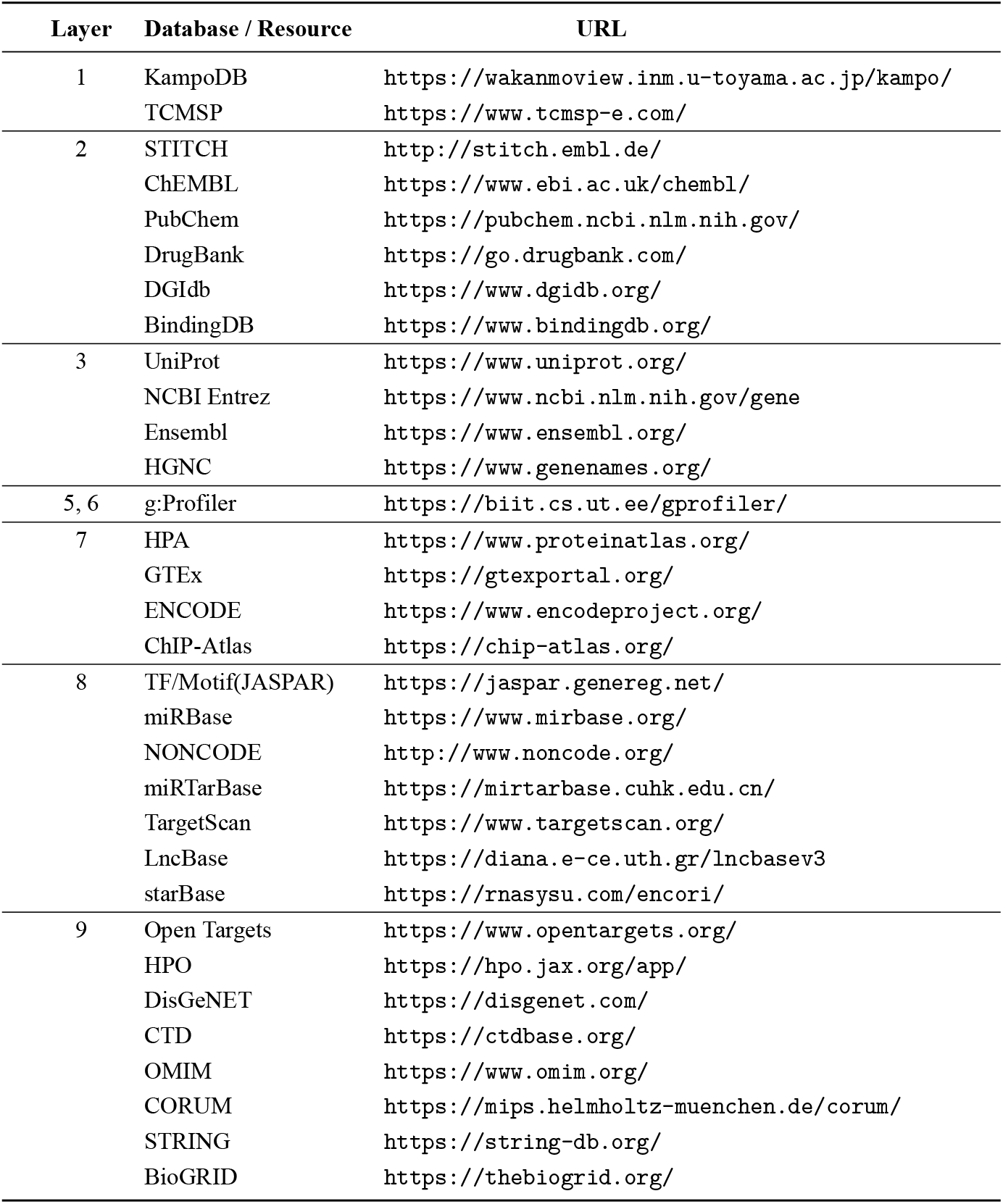
Data sources and implementation details for Phase 1 (Data-Driven Knowledge Construction).

**Table S1.**
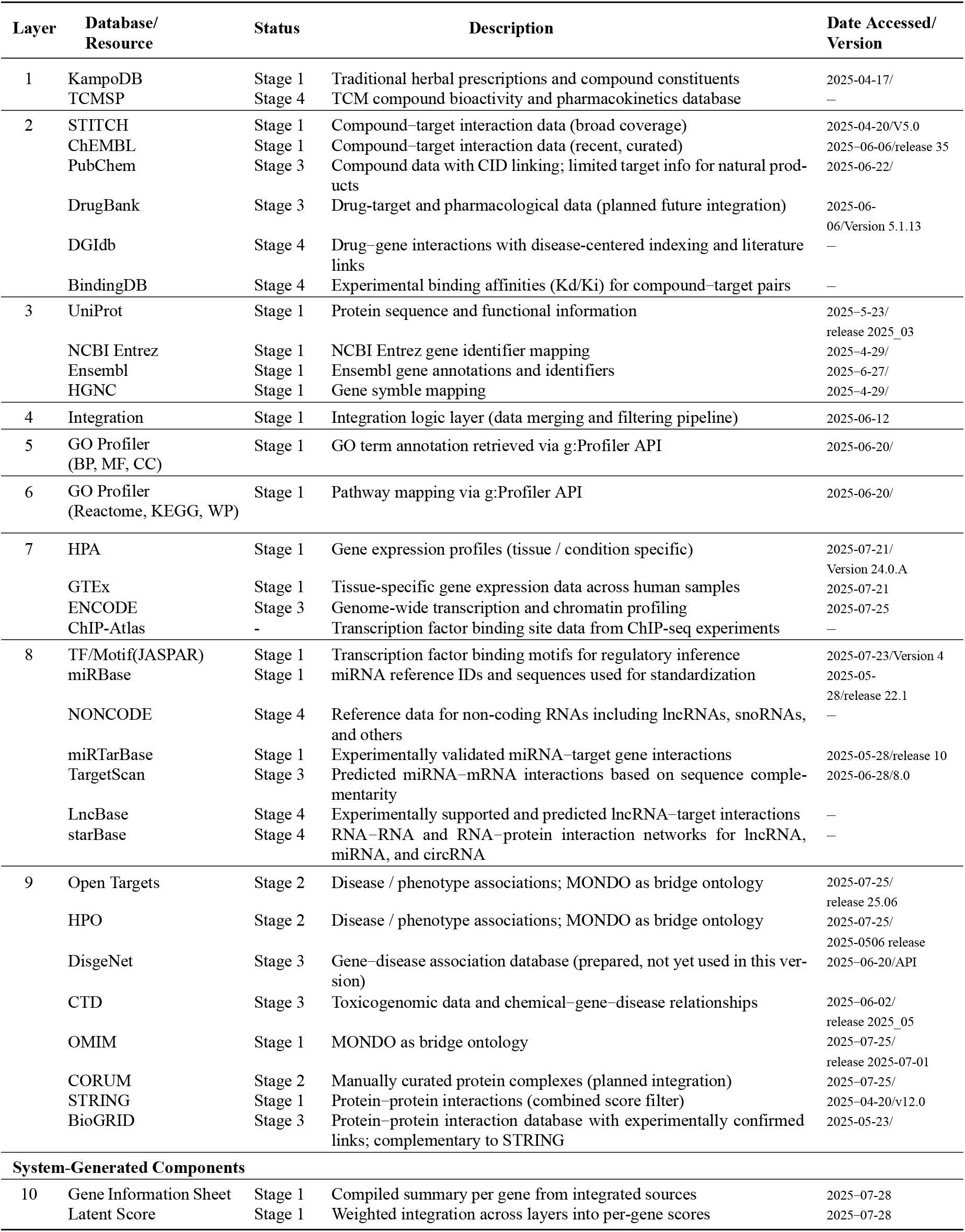
Data sources and implementation details for Phase 1 (Data-Driven Knowledge Construction). System-generated components developed in this study are listed separately at the bottom of the table.

### S2 Phase 2: Data Sources and Module Status for Hypothesis Generation

This section summarizes the modules and resources used in Phase 2 of the system, which focuses on hypothesis generation based on the integrated knowledge base constructed in Phase 1.

Phase 2 consists of eight conceptual layers, each corresponding to a functional module in the hypothesis-generation pipeline (see Figure 2B). These include semantic summarization (Character Sheets), knowledge graph construction, clustering, and multi-level hypothesis synthesis. While several components have been implemented at the prototype level, others remain in the planning stage.

The following table lists each layer, its purpose, representative resources or methods, and the current implementation status.

The classification follows four stages:

- **Stage 1**: Fully implemented in the present system
- **Stage 2**: Prototype implemented; requires further refinement
- **Stage 3**: Data and method prepared but not yet integrated
- **Stage 4**: Planned for future development

**Table S2.** Phase 2 modules for hypothesis generation: description, resources, and implementation status.

| Layer | Function / Resource | Implementation Status |
| --- | --- | --- |
| 1. Character Sheet | GPT-based semantic summarization (Template 1 → 2) | Stage 2: Prototype implemented (single model, extendable to multi-model) |
| 2. Knowledge Graph & Clustering | Triple conversion, Leiden clustering, UniProt, GO, Reactome, STRING | Stage 2: Graph and clustering implemented; pipeline in early use |
| 3. Semantic Screening | PubMedBERT, BioBERT, SciBERT (planned ensemble) | Stage 4: Planned module for semantic filtering and prioritization |
| 4. Biological Screening | Pathway-based logical layering and filtering | Stage 4: Planned integration prior to hypothesis synthesis |
| 5. Cluster-level Hypothesis | LLM-based hypothesis generation (Template 3 → 4) from selected CS | Stage 2: Implemented using GPT; designed for iterative prompting |
| 6. Formula-level Hypothesis | Aggregation of cluster-level hypotheses (Template 5 → 6) | Stage 2: Early implementation; cross-cluster synthesis logic applied |
| 7. Hypothesis Verification | Rule-based or LLM-based verification of mechanism | Stage 4: Planned validation logic (textual and symbolic reasoning) |
| 8. Visualization | Graph rendering and export for network-based hypothesis illustration | Stage 3: script implementation possible |

### S3 Phase 2: Cluster 14 Hypothesis (full text)

The following block presents the hypothesis exactly as generated by the system.

{

“cluster_id”: 14,

“master_hypothesis_title”: “Orchestrating Inflammatory Response via NFKBIA and TLR4: A Multi-Pathway Modulation Approach”,

“master_hypothesis_abstract”: “This hypothesis posits that the modulation of inflammatory responses in diseases such as psoriasis and depression is primarily orchestrated through the central role of NFKBIA and *TLR4* within the NF-κB and TLR signaling pathways. NFKBIA, ranked as the most topologically critical gene, acts as a key regulator by inhibiting the activity of the NF-κB complex, thereby preventing excessive inflammatory responses. TLR4 functions as an essential upstream sensor, activating downstream NF-κB signaling via its involvement in both NF-κB and HIF-1 pathways, thus tying immune activation to metabolic responses. The interaction of NFKBIA with traditional anti-inflammatory compounds like Kaempferol and Isoliquiritigenin highlights a therapeutic potential in modulating this pathway to alleviate inflammation-driven diseases. Furthermore, Paeoniflorin and Puerarin are hypothesized to modulate TLR4 activity, potentially attenuating the inflammatory responses associated with depression and atherosclerosis by dampening NF-κB and HIF-1 signaling. The combined modulation of these pathways by these compounds provides a comprehensive approach to managing inflammatory diseases by targeting multiple nodes within the network.”,

“key_players_and_roles”: [

{ “entity”: “Gene: NFKBIA”, “topological_rank”: 0, “role_in_hypothesis”: “Serves as the central regulator of the NF-κB pathway, inhibiting NF-κB activity to prevent excessive inflammation.” },

{ “entity”: “Gene: TLR4”, “topological_rank”: 1, “role_in_hypothesis”: “Functions as a critical upstream sensor and activator of NF-κB and HIF-1 pathways, linking immune and metabolic responses.” },

{ “entity”: “Gene: NFKB1”, “topological_rank”: 2, “role_in_hypothesis”: “Forms part of the NF-κB complex, a direct target for regulatory actions by NFKBIA, influencing inflammatory gene expression.” },

{ “entity”: “Gene: CHUK”, “topological_rank”: 3, “role_in_hypothesis”: “Facilitates the phosphorylation and subsequent degradation of IκB proteins, thus indirectly modulating NF-κB activity.” },

{ “entity”: “Gene: RELA”, “topological_rank”: 4, “role_in_hypothesis”: “A core component of the active NF-κB transcription factor complex, directly involved in inflammatory gene transcription.” },

{ “entity”: “Disease: Psoriasis”, “topological_rank”: “N/A”, “role_in_hypothesis”: “An inflammatory condition potentially mitigated by targeting NF-κB pathway components to reduce skin inflammation.” },

{ “entity”: “Disease: Depression”, “topological_rank”: “N/A”, “role_in_hypothesis”: “Linked to inflammatory processes modulated through NF-κB and TLR4 pathways, where targeted intervention may alleviate symptoms.” },

{ “entity”: “Disease: Ectodermal dysplasia with immunodeficiency”, “topological_rank”: “N/A”, “role_in_hypothesis”: “Characterized by immune dysregulation, potentially due to aberrant NF-κB pathway activity.” },

{ “entity”: “Compound: Paeoniflorin”, “topological_rank”: “N/A”, “role_in_hypothesis”: “Proposed to modulate TLR4 activity, potentially reducing NF-κB pathway activation and inflammatory responses.” },

{ “entity”: “Compound: Puerarin”, “topological_rank”: “N/A”, “role_in_hypothesis”: “Likely acts on TLR4 to modulate both NF-κB and HIF-1 pathways, offering a multi-faceted approach to inflammation control.” }

],

“proposed_validation_strategy”: “To validate this hypothesis, a multi-tiered experimental approach is proposed. First, co-immuno-precipitation assays should be conducted to confirm the interactions between NFKBIA and NF-κB complex components, such as NFKB1 and RELA. Secondly, TLR4 activation assays using Paeoniflorin and Puerarin should be performed to evaluate their effects on NF-κB and HIF-1 pathway signaling. Additionally, conducting phosphorylation assays on CHUK can elucidate its role in IκB degradation. Finally, in vivo models of psoriasis and depression should be utilized to assess the therapeutic efficacy of these compounds in reducing inflammatory markers and symptoms.”

}

### S4 Phase 2: Final Hypothesis (full text)

The following block presents the final integrated hypothesis as generated by the system.

~~~
# Syakuyaku-Kanzo-to (芍薬甘草湯)
---
## Integrative Modulation of Multi-Pathway Networks for Comprehensive Disease Management
~~~

**Core Principle:**

Leveraging traditional compounds to modulate interconnected biological networks across inflammation, neuro-psychiatric, metabolic, and oncogenic pathways for holistic therapeutic outcomes.

**Abstract:**

This hypothesis proposes a comprehensive approach to disease management by strategically modulating key biological networks through traditional compounds. The formula Syakuyaku-Kanzo-to (芍薬甘草湯) integrates compounds such as Genistein, Melatonin, and Glycyrrhiza to target multiple pathways involved in inflammation, immune modulation, neuro-psychiatric regulation, metabolic balance, and anti-tumor activity. By influencing critical nodes such as IL6, MTOR, and CYP3A4, these compounds offer a multi-target strategy that addresses the underlying mechanisms of complex diseases like cancer, metabolic disorders, and neurodegenerative conditions. This network pharmacology approach not only enhances our understanding of disease pathogenesis but also paves the way for innovative treatments that align traditional wisdom with modern scientific insights.

~~~
---
## Four Functional Modules
### Inflammation & Immune Modulation
**Summary:** This module focuses on modulating inflammatory responses and immune pathways through key cytokines and signaling molecules.
- **Key Compounds:** Genistein, Melatonin, Glycyrrhiza, 3’,4’,7-Trihydroxyisoflavone
- **Key Gene Targets:** IL6, TNFRSF1A, CXCR4, MIF
- **Concrete Examples:**
 - Genistein modulates the IL6-TNFRSF1A axis to reduce inflammation and tumorigenic signaling (from Cluster 17).
 - Glycyrrhiza and 3’,4’,7-Trihydroxyisoflavone target the CXCR4-MIF axis to stabilize immune homeostasis (from Cluster 13).
### Neuro-Psychiatric Modulation
**Summary:** This module targets neurotransmitter pathways to regulate mood disorders and neuro-developmental conditions.
- **Key Compounds:** Serotonin, Daidzein, Glycyrrhiza
- **Key Gene Targets:** SLC6A4, HTR1A, GABRG2, MTOR
- **Concrete Examples:**
 - Serotonin and Glycyrrhiza modulate SLC6A4 and HTR1A to influence mood regulation in major depressive disorder (from Cluster 23).
 - Daidzein affects GABRG2 activity, offering therapeutic potential in anxiety and epilepsy (from Cluster 23).
### Metabolic & Endocrine Modulation
**Summary:** This module addresses metabolic disorders by influencing key hormonal and enzymatic pathways.
- **Key Compounds:** Genistein, Glycyrrhiza, Melatonin
- **Key Gene Targets:** CYP3A4, PTH1R, CYP24A1
- **Concrete Examples:**
 - Genistein modulates CYP3A4 activity to influence drug metabolism and viral infection progression (from Cluster 31).
 - Glycyrrhiza and Genistein enhance PTH1R signaling to improve bone density and calcium homeostasis (from Cluster 33).
### Anti-Tumor & Cell Proliferation
**Summary:** This module targets oncogenic pathways to inhibit tumor growth and promote apoptosis.
- **Key Compounds:** Genistein, Melatonin, Liquiritigenin
- **Key Gene Targets:** RET, PIK3CA, MTOR, PSEN1
- **Concrete Examples:**
 - Genistein inhibits RET-mediated pathways to reduce cancer cell proliferation in thyroid and lung cancers (from Cluster 34).
 - Melatonin modulates MTOR activity to rebalance growth signals in neurodevelopmental disorders and cancer (from Cluster 27).
---
## Synergy and Crosstalk
~~~

The modules interact through shared compounds like Genistein and Melatonin, which modulate multiple pathways such as inflammation, metabolic regulation, and tumor suppression. For instance, Genistein’s role in both the CYP3A4-centered metabolic network and the RET-mediated oncogenic pathways illustrates its broad therapeutic potential. Melatonin’s influence on both circadian rhythms and oxidative stress further exemplifies the interconnected nature of these biological systems.

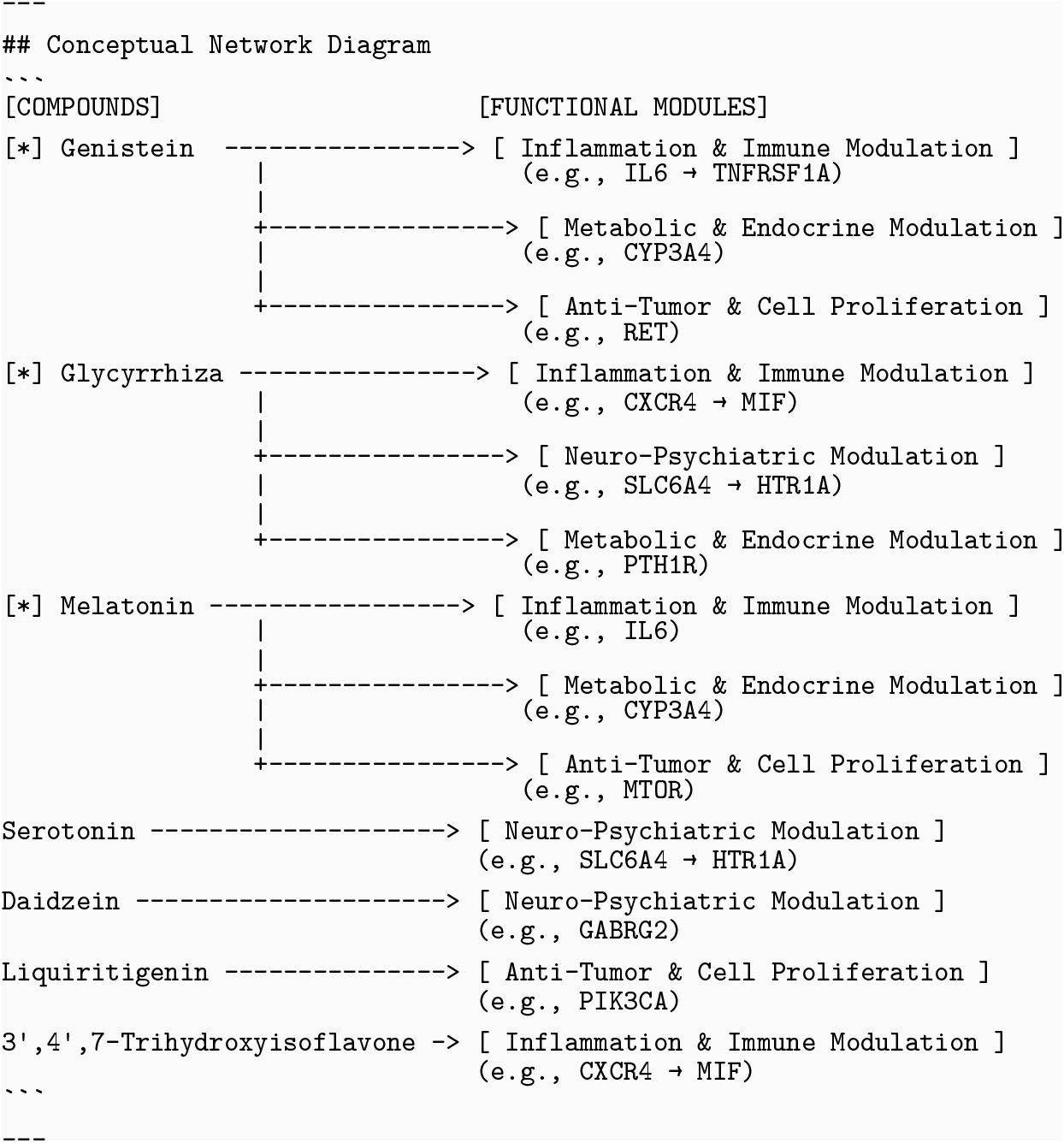

\## Concluding Vision

This integrative approach highlights the potential of traditional compounds to modulate complex disease networks, offering a holistic strategy for managing multifaceted conditions. By bridging traditional medicine with modern pharmacology, this hypothesis not only enhances therapeutic efficacy but also opens new avenues for personalized medicine, ultimately improving patient outcomes across a spectrum of diseases.

### S5 Manual validation of Cluster 14

This section provides the results of manual validation for the hypotheses generated in Cluster 14. Each subject–relation–object triple from the network (Figure 4) was validated through PubMed searches. For every relationship, at least one supporting reference (PMID and DOI) was identified, and a brief note was added to clarify the context. These validations confirm that the Cluster 14 hypothesis aligns with existing biomedical literature.

---

**Table S5.** Manual PubMed validation of relations depicted in Figure 4 (Cluster 14). Disease associations are labeled as corresponding to Figure 4 diseases.

| Subject | Relation | Object | Evidence (PMID / DOI ) | Year | Note |
| --- | --- | --- | --- | --- | --- |
| Paeoniflorin | inhibits | *TLR4* | PMID: 29740218/<br>DOI:10.2147/CMAR.S160292<br>PMID: 38331105<br>DOI:10.1016/j.taap.2024.116846 | 2018<br>2024 | **Paeoniflorin** (Pae) inhibits *TLR4*. A 2018 glioblastoma study showed Pae promotes ubiquitin–proteasome–mediated degradation of *TLR4*, suppressing NF-κB. A 2024 cachexia study confirmed Pae inhibits the ***TLR4*/NF-κB** axis and ameliorates muscle atrophy in vitro and in vivo. |
| Puerarin | inhibits | *TLR4* | PMID: 38905846/<br>DOI:10.1016/j.phymed.2024.155813 | 2024 | “**Puerarin** bound to Toll/interleukin-1 receptor (TIR) domain of MyD88 protein, hindering its binding with ***TLR4***, ultimately resulting in downstream NF-κB p65 and JNK/FoxO1 signaling inactivation.” |
| Kaempferol | activates | *NFKBIA* | PMID: 17278014/<br>DOI:10.1007/s10522-007-9083-9 | 2007 | **Kaempferol** upregulates IκBα (*NFKBIA*), preventing NF-κB nuclear translocation and reducing inflammation. |
| Isoliquiritigenin | activates | *NFKBIA* | PMID: 18295200/<br>DOI:10.1016/j.ejphar.2008.01.032 | 2008 | **Isoliquiritigenin** (ILG) stabilizes IκBα (*NFKBIA*) by inhibiting its phosphorylation and degradation, thereby preventing NF-κB nuclear translocation. |
| *TLR4* | activates | NF-κB | PMID: 29740218/<br>DOI:10.2147/CMAR.S160292<br>PMID: 38905846/<br>DOI:10.1016/j.phymed.2024.155813 | 2018<br>2024 | A 2018 study described **NF-κB** as a canonical downstream effector of *TLR4*. A 2024 study showed this signaling was suppressed by puerarin. |
| *TLR4* | activates | HIF-1 | PMID: 39006190/<br>DOI:10.2147/DDDT.S436359 | 2024 | In the cited study, this refers specifically to HIF-1α, the regulatory subunit of the **HIF-1** complex. UC (DSS) model; berberine suppresses the ***TLR4*/NF-κB/HIF-1α** axis. Counts as indirect (via NF-κB) pathway-level evidence. |
| *NFKBIA* | inhibits | NF-κB | PMID: 28428267/<br>DOI:10.1530/ERC-17-0033 | 2017 | *NFKBIA* functions as inhibitor of **NF-κB** signaling; genetic variant linked to altered inflammatory response. |
| *NFKB1*<br>*RELA* | is subunit of | NF-κB | PMID: 39337282/<br>DOI: 10.3390/ijms25189793 | 2024 | *NFKB1* (p105/p50) and *RELA* (p65) are canonical **NF-κB** subunits. |
| *CHUK* | inhibits | *NFKBIA* | PMID: 39622883/<br>DOI:10.1038/s41598-024-80101-1 | 2024 | **CHUK** phosphorylates IκBα (*NFKBIA*), leading to its degradation and relieving NF-κB inhibition. |
| NF-κB | associated with | Psoriasis | PMID: 34460100/<br>DOI:10.1111/bph.15673 | 2021 | NF-κB inhibition (e.g., by H-151) reduced cytokines and improved psoriatic lesions, indicating NF-κB activation contributes to pathology (**corresponding to Figure 4 disease**). |
| NF-κB | associated with | Depression | PMID: 37032645/<br>DOI:10.1111/cns.14207 | 2023 | In CUMS-exposed rats, NF-κB p65 activation was upregulated, contributing to **depression-like** behaviors. TaVNS suppressed NF-κB signaling, reducing neuroinflammation and alleviating depressive phenotypes. |
| NF-κB | associated with | Ectodermal dysplasia with immunodeficiency | PMID: 11242109/<br>DOI:10.1038/85837 | 2001 | Mutations in IKBKG (NEMO), the regulatory subunit of the IKK complex, impair NF-κB signaling. This impairment underlies **ectodermal dysplasia with immunodeficiency** (EDA-ID) and related syndromes, linking defective NF-κB activity to the disease phenotype. |
| *TLR4* | associated with | Depression | PMID: 32964428/<br>DOI:10.1111/bph.15261 | 2020 | Arctigenin suppressed HMGB1/*TLR4*/NF-κB signaling, reduced **depressive-like behaviors** (TST, FST). |
| HIF-1 | associated with | Depression | PMID: 36944982/<br>DOI: 10.1186/s12974-023-02765-2 | 2023 | In the cited study, this refers specifically to HIF-1α, measured as the disease-associated factor underlying **depression-like** behaviors. |

